# Human *RAP2A* Homolog of the *Drosophila* Asymmetric Cell Division Regulator *Rap2l* Targets the Stemness of Glioblastoma Stem Cells

**DOI:** 10.1101/2025.01.28.635292

**Authors:** Maribel Franco, Ricardo Gargini, Víctor M. Barberá, Daniel Becerra, Miguel Saceda, Ana Carmena

## Abstract

Asymmetric cell division (ACD) is a fundamental process to balance cell proliferation and differentiation during development and in the adult. Cancer stem cells (CSCs), a very small but highly malignant population within many human tumors, are able to provide differentiated progeny by ACD that contribute to the intratumoral heterogeneity, as well as to proliferate without control by symmetric, self-renewing divisions. Thus, ACD dysregulation in CSCs could trigger cancer progression. Here we consistently find low expression levels of *RAP2A*, the human homolog of the *Drosophila* ACD regulator *Rap2l*, in glioblastoma (GBM) patient samples, and observe that scarce levels of *RAP2A* are associated with poor clinical prognosis in GBM. Additionally, we show that restitution of RAP2A in GBM neurosphere cultures increases the ACD of glioblastoma stem cells (GSCs), decreasing their proliferation and expression of stem cell markers. Our results support that ACD failures in GSCs increases their spread, and that ACD amendment could contribute to reduce the expansion of GBM.

## Introduction

Asymmetric cell division (ACD) is a universal mechanism for generating cellular diversity during development as well as for regulating tissue homeostasis in the adult (1). Stem and progenitor cells undergo ACD to simultaneously give rise to a self-renewing stem/progenitor cell and to a daughter cell that is committed to enter a differentiation program. Remarkably, over the past years it has become apparent that undermining this critical process can lead to tumor-like overgrowth (2). This link between failures in the process of ACD and tumorigenesis was first established using the *Drosophila* neural stem cells, called neuroblasts (NBs), as a model system (3). In this work, authors showed how *Drosophila* larval mutant brains for ACD regulatory genes were able to induce the formation of massive tumoral masses after weeks of being transplanted into the abdomen of wild-type fly hosts. Intriguingly, some genes firstly identified in *Drosophila* as tumor suppressor genes, such as *discs large 1* (*dlg1*), *lethal (2) giant larvae* (*l(2)gl*) and *brain tumor* (*brat*) (4), were later uncovered as ACD regulators (5-11), further supporting the connection between compromised ACD and tumorigenesis.

Tumor initiating cells or cancer stem cells (CSCs) were originally identified in acute myeloid leukemia (AML) (12,13) but, to date, several studies have also revealed the existence of CSCs in multiple solid tumors (14). These CSCs represent a very small but highly malignant population within the tumor, able to proliferate without control by symmetrical, self-renewal divisions, as well as to provide differentiated progeny by ACD that contribute to the heterogeneity of the tumor. These properties, along with their quiescent state, increase in metabolic activity and high capacity of DNA repair, make CSCs extremely resistant to radio- and chemotherapy and responsible for malignant relapse (14-18). Thus, understanding the unique nature of these cells and their landmarks is crucial to target CSCs and hence to completely destroy the tumor. An intriguing possibility is that an imbalance in the mode of CSC divisions, favoring symmetric renewal divisions in detriment of asymmetric divisions, might contribute to the transition from a chronic (low grade) to an acute (high grade) phase in the cancer progression (19,20).

Glioblastoma (GBM), the most aggressive and lethal brain tumor with very poor prognosis, is among the solid tumors in which the presence of CSCs, called glioma or glioblastoma stem cells (GSCs), has been proven. As other CSCs, GSCs are responsible for the formation, maintenance, resistance to conventional therapy and consequent relapse of GBM (21-25). Several GSC biomarkers, such as CD133 and Nestin, and signaling pathways, such as Notch and Hedgehog, have been identified to help isolate and target these crucial cells (14,26,27). Despite of that, there are still many unknowns regarding the nature, properties and origin of CSCs in general and of GSCs in particular.

Here, we analyze the consequences of compromising ACD in the GSCs within GBM neurosphere cell cultures. Specifically, we show that *RAP2A*, the human homolog of *Rap2l*, which encodes a *Drosophila* small GTPase and novel ACD regulator, displays low expression levels in GBM, and that restitution of RAP2A in GBM neurosphere cultures increases the ACD and decreases the expression of stem cell markers in the GSCs. Our results support that failures in ACD increase GSC proliferation promoting their spreading and that ACD amendment might contribute to reduce the expansion of GBM.

## Results

### RAP2A is highly downregulated in GBM patients

As a first approach to analyze whether ACD might be impaired in GBM, we analyzed the expression levels of human homologs of 21 *Drosophila* ACD regulators in a GBM microarray (28) (**Fig. 1A**). We also investigated which of those genes were more similar in expression pattern to the others by performing a Gene Distance Matrix (**Fig. 1B**). The human genes *TRIM2* and *RAP2A* showed the lowest levels in the GBM patient samples analyzed. *TRIM2* belongs to the same family that *TRIM3* and *TRIM32*, all of them related to the *Drosophila* ACD regulator gene *brat. TRIM3*, the closest homolog of *brat*, has been previously found to be expressed at low levels in GBM (29). Hence, we concentrated on *RAP2A*, the second human gene consistently found at very low levels in all the patient samples (**Fig. 1A and Fig. S1**). *RAP2A* is also known to have a tumor suppressor role in the context of glioma migration and invasion (30,31). First, to further support the microarray data expression levels in a bigger sample size, we performed *in silico* analyses taking advantage of established datasets of Glioblastoma IDH wild-type, such as “The Cancer Genome Atlas” (TCGA) and “Gravendeel” cohorts. We confirmed that low *RAP2A* expression levels in human GBM are associated with a poor prognosis (**Fig. 1C**). In a multivariate analysis in the TCGA cohort, when we included the variables RAP2A expression, patient age and standard chemo/radiotherapy treatment in the model, it was observed that RAP2A levels and patient age have a significant implication in patient survival as an independent variable. In fact, specific values were significant in predicting overall survival (OS). For example, “RAP2A expression” with a hazard ratio (HR) of 0.866 (95% CI 0.760-0.986) and a *p*-value of 0.03, and “patient age” with a hazard ratio (HR) of 1.030 (95% CI 1.022-1.037) and a *p*-value of <0.001. Conversely, standard chemo/radiotherapy treatment did not remain as an independent variable, with a hazard ratio (HR) of 0.820 and a *p*-value of 0.053. We also analyzed RAP2A levels in the different GBM subtypes (proneural, mesenchymal and classical) in the TCGA cohort and their prognostic relevance. The proneural subtype that displayed RAP2A levels significantly higher than the others was the subtype that also showed better prognosis (**Fig. S2**). Hence, we focused on the human gene *RAP2A* to analyze its potential role in ACD and its relevance in GBM.

**Figure 1.**
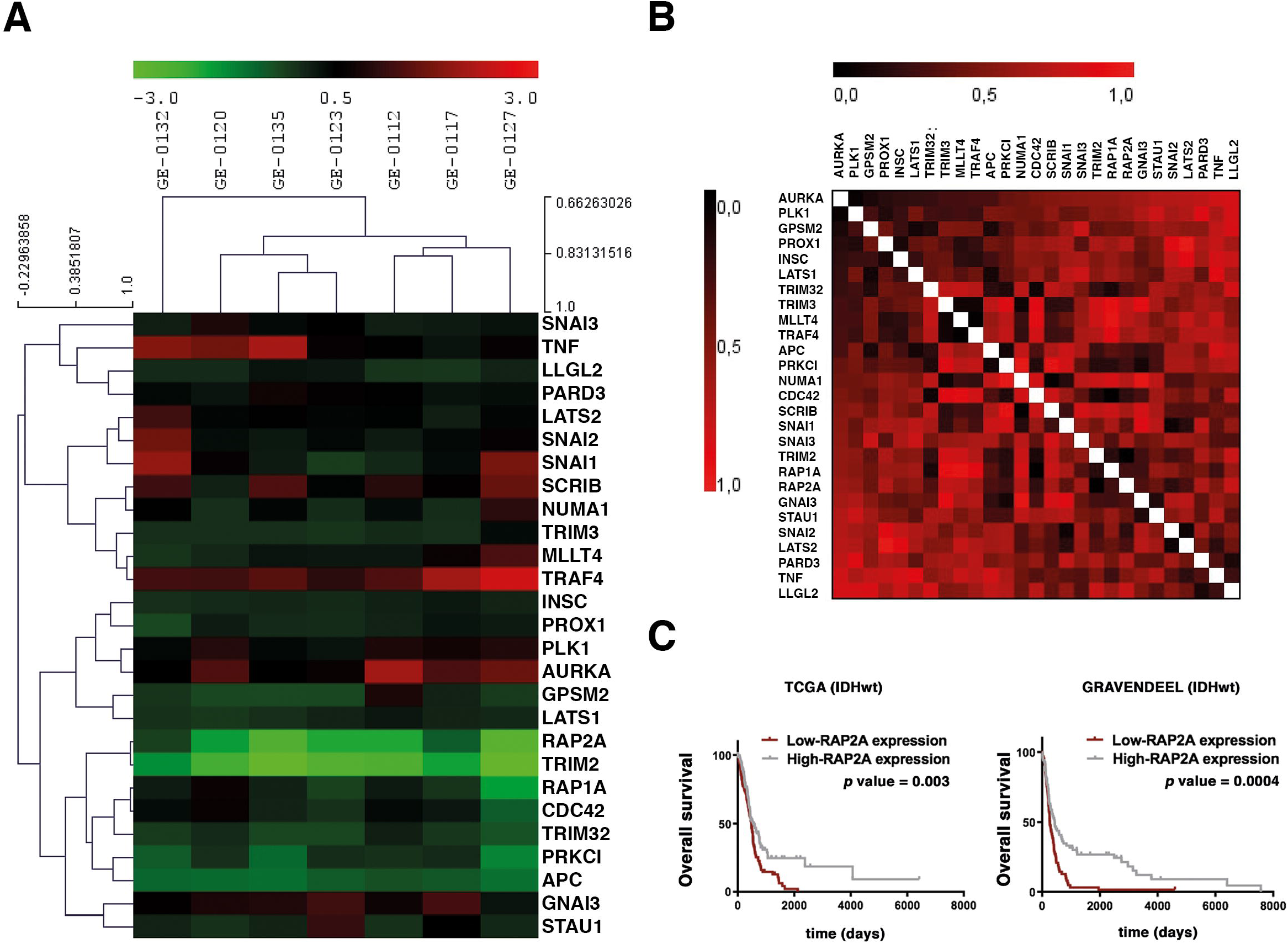
RAP2A is highly downregulated in GBM patients. (**A, B)** Analysis of GBM patient versus control samples focused on the level of expression of human homologs of *Drosophila* ACD regulators by Hierarchical clustering (**A**) and the Gene Distance Matrix (**B**). Color-coded scale bar indicates fold level of expression. **(C)** Kaplan-Meier survival curves corresponding to patients in the TCGA (IDHwt) and Gravendeel (IDHwt) cohort in GBM datasets. Low *RAP2A* expression levels in human GBM are associated with a poor prognosis.

### Drosophila RAP2A homolog Rap2l regulates ACD

We previously showed that the *Drosophila* gene *Rap1* regulates the ACD process (32). The closest human homolog of *Drosophila Rap1* is *RAP1A*, while human *RAP2A* is more similar to *Drosophila Rap2l*, which has not been previously analyzed in the context of ACD. Hence, we first wanted to investigate whether *Rap2l* was, like *Rap1*, implicated in ACD regulation. With that aim, we started analyzing the ACD of neural stem cells (neuroblasts, NBs) and intermediate neural progenitors (INPs) within *Drosophila* larval brain type II NB lineages (NBII) (**Fig. 2A**) (8,33,34). In NBII lineages, there is one NB that expresses the transcription factor Deadpan (Dpn) and several INPs, which express both Dpn and the transcription factor Asense (Ase) (**Fig. 2A**). We first observed the presence of ectopic NBs (eNBs; Dpn^+^ Ase^-^) after downregulating *Rap2l* in NBII lineages in 4 out of 5 mutant brains analyzed (n = 34 NB lineages), while in control lineages only one NB was usually found (**Fig. 2B**). To further determine the specificity and penetrance of the phenotype, we repeated this experiment analyzing this and two additional *Rap2l* RNAi lines, substantially increasing the number of the samples (both the number of NB lineages and the number of brains analyzed) (**Fig. S3**). All the different lines showed a highly significant number of ectopic NBs (**** *p* < 0.001; n> 120 NB lineages analyzed) in all or almost all of the different brains analyzed (n>14) (see **Fig. S3** for details). This result suggested that the process of ACD was impaired within those *Rap2l* mutant NBII lineages. To more directly support that, we analyzed the localization of key ACD regulators in metaphase NBs and INPs of NBII lineages, such as the apical proteins aPKC and Canoe (Cno), and the cell-fate determinant Numb (35-37). In control metaphase NBs and INPs, those ACD regulators form crescents at the apical (aPKC and Cno) or basal (Numb) poles of the dividing cell (**Fig. 2C**). The apical localization of aPKC was not significantly altered after downregulating *Rap2l* in NBII lineages; however, the localization of Cno and Numb did fail in most of the *Rap2l* mutant brains analyzed (in 5 out of 6 in the case of Cno, and in 3 out of 4 in the case of Numb) (**Fig. 2C**). All these results strongly suggested that Rap2l, as Rap1, was an ACD regulator. Hence, we came back to the human homolog of *Drosophila Rap2l, RAP2A*, to determine its potential relevance in the context of the ACD of GSCs.

**Figure 2.**
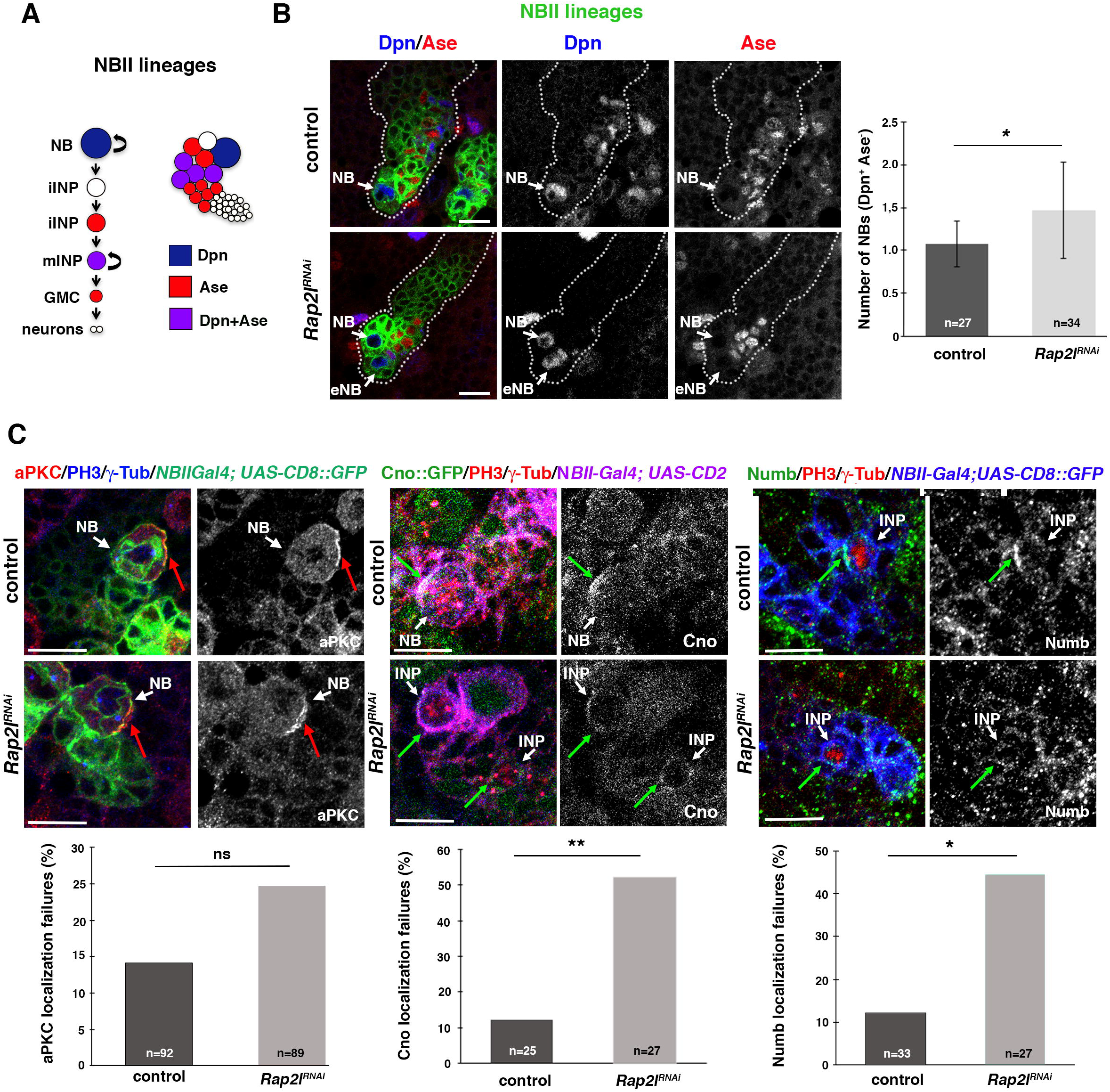
*Drosophila RAP2A* homolog *Rap2l* regulates ACD. (**A**) *Drosophila* NBII lineage; the only NB per lineage expresses the transcription factor Dpn, while mature INPs express both Dpn and Ase; iINP, immature INP; mINP, mature INP; GMC, Ganglion Mother Cell. **(B)** Confocal micrographs showing a control larval brain NBII lineage with one NB (Dpn^+^ Ase^-^) and an NBII lineage in which *Rap2l* has been downregulated displaying an ectopic NB (eNB). **(C)** *Rap2l* downregulation in NBII lineages by the *wor-Gal4 ase-Gal80* (an NBII-specific driver) causes Cno and Numb localization failures (green arrows) in dividing progenitors (NB or INPs), while aPKC localization is not significantly altered (red arrows). Data shown in the scaled bar graphs was analyzed with a U Mann-Whitney test (B) or a Chi-square test (C), \**p*<0.05, \*\**p*<0.01, ns, not significant, n=number of NB lineages analyzed (in B) or number of dividing cells analyzed (in C); scale bar: 10 μm.

### RAP2A expression in GBM neurosphere cultures reduces the stem cell population

After confirming by *in silico* analyses that low *RAP2A* expression levels in GBM patients were associated to a poor prognosis, we decided to use established GBM cell lines to investigate in neurosphere cell cultures whether the downregulation of *RAP2A* was affecting the properties of GSCs. We tested six different GBM cell lines finding similar mRNA levels of *RAP2A* in all of them, and significantly lower levels than in control Astros (**Fig. 3A**). We focused on the GBM cell line called GB5, which grew well in neurosphere cell culture conditions, for further analyses. Given that stem cell markers, such as CD133, SOX2 and NESTIN are landmarks of GSCs, we started analyzing their expression after restoring RAP2A in the neurosphere culture (**Fig. 3B**). Immunofluorescences of each of those markers revealed a significant reduction in the protein intensity in the RAP2A-expressing neurospheres (GB5-RAP2A) compared to the control neurospheres (GB5) (**Fig. 3C**). Likewise, RT-PCRs and Westerns blots also showed a significant decrease in the mRNA and protein expression levels, respectively, of those stem cell markers (**Fig. 4**). Hence, RAP2A seems to be contributing to decrease the stem cell population within GBM neurosphere cultures.

**Figure 3.**
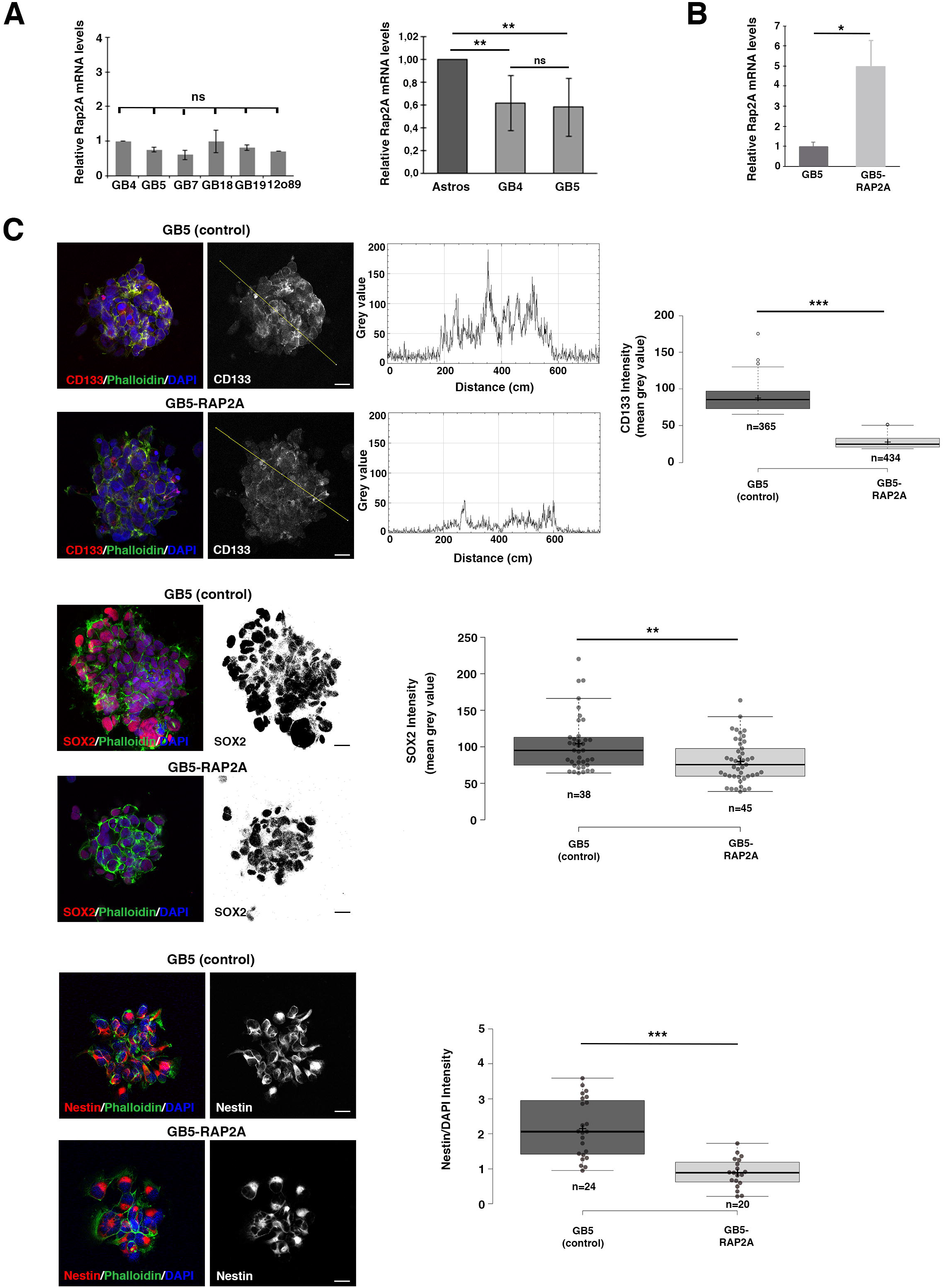
*RAP2A* expression in GBM neurosphere cultures reduces the stem cell population. **(A)** Different GBM cell lines show similar *RAP2A* mRNA levels and significantly lower levels than in control Astros. **(B)** *RAP2A* mRNA levels are significantly higher in the GB5 line after infecting this line with *RAP2A* (GB5-RAP2A). Data shown in the scaled bar graphs in A and B was analyzed with an ANOVA and a T-test, respectively; error bars show the SD; n= 2 (in A) and 3 (in B) different experiments. **(C)** Immunofluorescences of the GSC stem cell markers CD133, SOX2 and Nestin reveal a significant reduction in the protein intensity in the GB5 RAP2A-expressing neurospheres (GB5-RAP2A) compared to the control neurospheres (GB5). Data shown in the box plots was analyzed with a U Mann-Whitney for CD133 and Nestin and with a T-test for Sox; the central lines represent the median and the box limits the lower and upper quartiles, as determined by R software; crosses represent sample means; error bars indicate the SEM; n= number of sample points; *p* values: \**p*<0.05, \*\**p*<0.01, \*\*\**p*<0.001, ns, not significant; scale bar: 10 μm.

**Figure 4.**
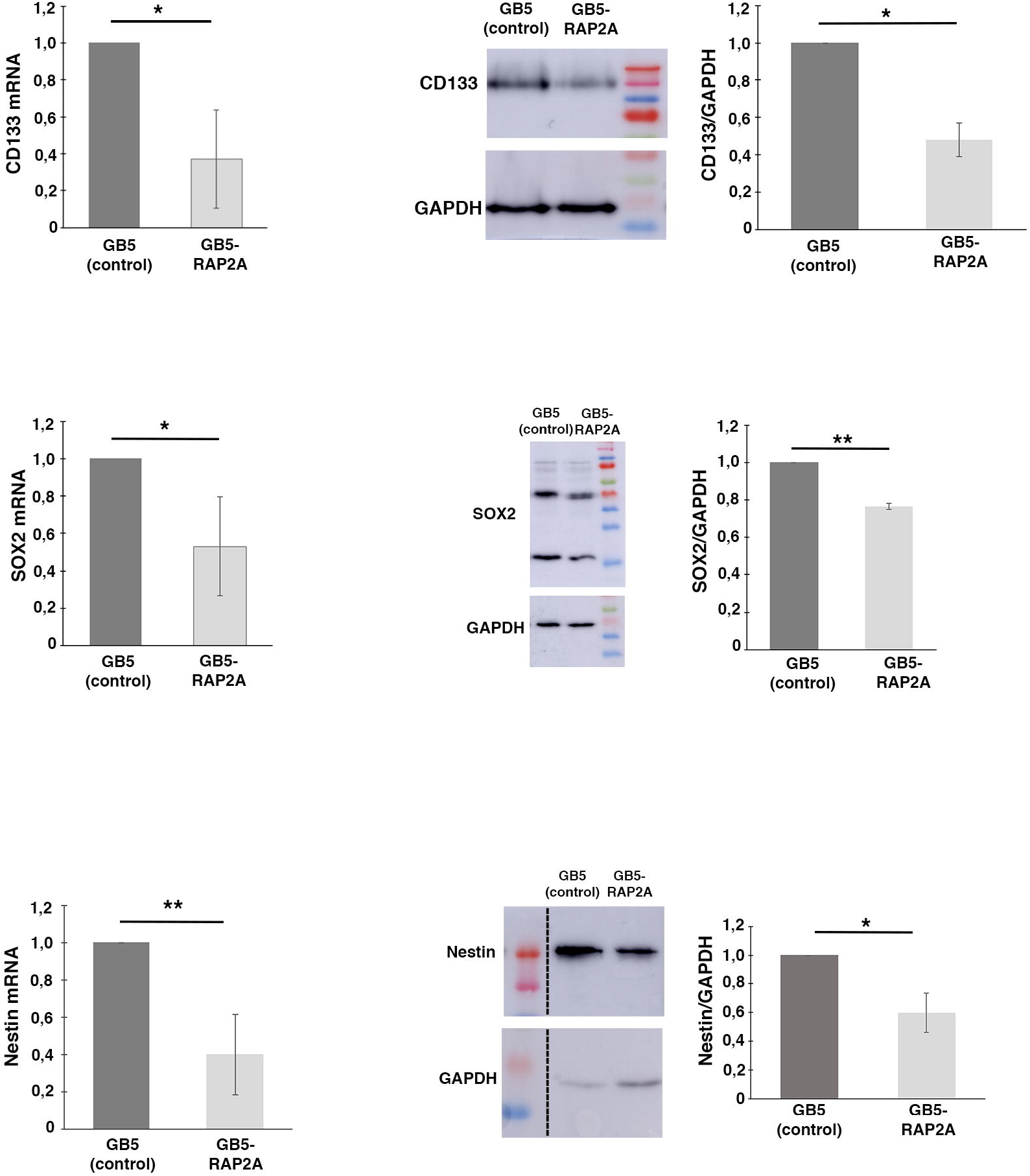
*RAP2A* expression in GBM neurosphere cultures reduces the stem cell population. RT-PCRs and Westerns blots show a significant decrease in the mRNA and protein expression levels, respectively, of the GSC markers CD133, SOX2 and Nestin in the GB5 RAP2A-expressing neurospheres (GB5-RAP2A) compared to the control neurospheres (GB5). Data shown in the scaled bar graphs was analyzed with an unpaired two-tailed Student’s *t* test; error bars show the SD; n= 3 different experiments; *p* values: \**p*<0.05, \*\**p*<0.01.

### RAP2A expression in GBM neurosphere cultures decreases cell proliferation and sphere size

Next, we wondered whether the reduction in the stem cell population after expressing *RAP2A* in neurosphere cultures was reflected in the level of cell proliferation and, consequently, in the size of the neurospheres. To investigate this, we first analyzed the expression of the proliferation marker and key tool in cancer diagnostics Ki-67, highly expressed in cycling cells but absent in quiescent (G0) cells (38,39). We detected a significant decrease in the number of proliferating cells per neurosphere in those cultures expressing *RAP2A* (**Fig. 5A**). Moreover, the average area size of these GB5-RAP2A neurospheres was also significantly reduced compared to control GB5 neurospheres (**Fig. 5B**). Thus, RAP2A promotes a decrease in cell proliferation.

**Figure 5.**
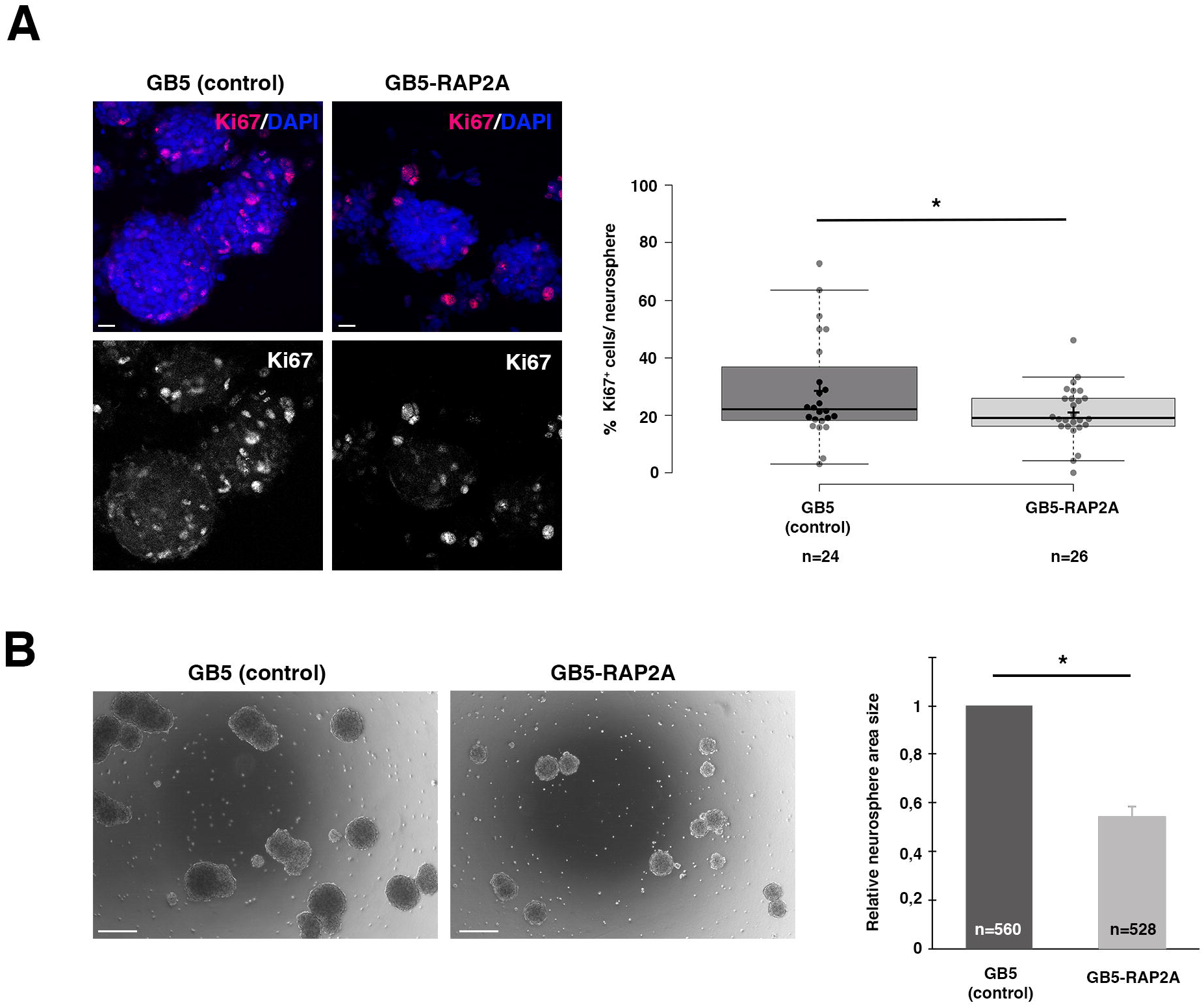
*RAP2A* expression in GBM neurosphere cultures decreases cell proliferation and sphere size. **(A)** GB5 neurospheres expressing RAP2A (GB5-RAP2A) show a significantly lower number of Ki67 expressing cells per neurosphere than control GB5 neurospheres. Data shown in the box plots was analyzed with a T-test; the central lines represent the median and the box limits the lower and upper quartiles, as determined by R software; crosses represent sample means; error bars indicate the SEM; n= number of sample points. **(B)** GB5 neurospheres expressing RAP2A (GB5-RAP2A) show a significant decrease in their size compared to control GB5 neurospheres of the same stage. Data shown in the scaled bar graphs was analyzed with an unpaired two-tailed Student’s T-test; error bars show the SD; n= total number of neurospheres of 3 different experiments; *p* values: \**p*<0.05; scale bars: 10 μm (in A) and 100μm (in B).

### RAP2A expression in GBM neurosphere cultures fosters ACD in GSCs

Given the reduction in cell proliferation and in the stem cell population after adding RAP2A to the control GB5 neurosphere cultures, we wanted to investigate whether RAP2A was favoring an asymmetric mode of cell division in these cells. Thus, we decided to follow over time the GB5-RAP2A and the control GB5 neurospheres, analyzing the number of cells per neurosphere. We reasoned that an even number of cells per neurosphere would be indicative of symmetric cell divisions, while an odd number of cells would point to an asymmetric mode of cell division (see also Material and methods for details). We observed a significant increase of neurospheres with an odd number of cells in the GB5-RAP2A neurospheres (n= 124 neurospheres analyzed) compared to control GB5 neurospheres (n=153 neurospheres) (**Fig. 6A**). This result was very suggestive of an increase in the number of ACDs in GB5-RAP2A neurosphere cells. To further support this result, we decided to analyze the distribution of the cell fate determinant NUMB, a key conserved ACD regulator whose asymmetric distribution in the mother and progeny ultimately promotes the repression of stem cell self-renewal in the daughter cell in which is segregated (40-44). We observed a significant increase in the number of GB5-RAP2A dividing cells that showed an asymmetric localization of NUMB in the progeny, compared to control GB5 dividing cells (**Fig. 6B**). Thus, all these results strongly supported a function of RAP2A promoting ACD in the GSCs.

**Figure 6.**
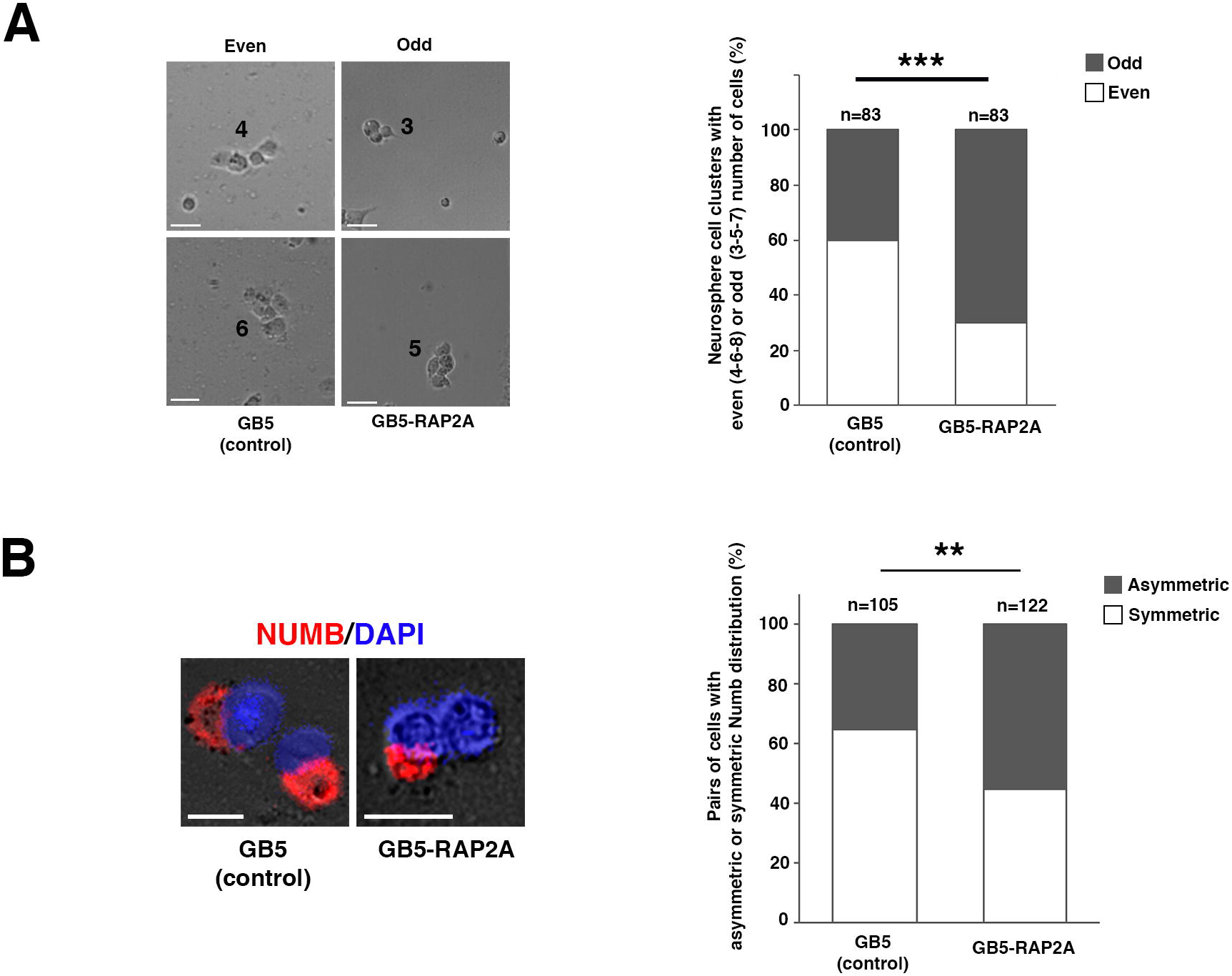
*RAP2A* expression in GBM neurosphere cultures fosters ACD in GSCs. **(A)** Early stage GB5 neurospheres expressing RAP2A (GB5-RAP2A) show an odd number of cells (3-5-7) significantly more frequently than control GB5 neurospheres, which show more frequently an even number of cells (4-6-8). **(B)** GB5-RAP2A dividing cells show a significant increase in the number of asymmetric NUMB localization in the progeny, compared to control GB5 dividing cells. Data shown in the bar graphs was analyzed with a Chi-Square test with Yates correction; n= number of neurosphere cell clusters analyzed. *p* values: \*\**p*<0.01; \*\*\**p*<0.001scale bar: 20 μm.

## Discussion

ACD is an evolutionary conserved mechanism used by stem and progenitor cells to generate cell diversity during development and to regulate tissue homeostasis in the adult. CSCs, present in many human tumors, can divide asymmetrically to generate intratumoral heterogeneity or symmetrically to expand the tumor by self-renewal. Over the past 15 years, it has been suggested that dysregulation of the balance between symmetric and asymmetric cell divisions in CSCs, favoring symmetric divisions, can trigger tumor progression in different types of cancer (19,20,45) including mammary tumors (43,46), glioblastoma (29), oligodendrogliomas (47,48), colorectal cancer (49,50) and hepatocellular carcinoma (50). Thus, it is of great relevance to get a deeper insight into the network of regulators that control ACD, as well as the mechanisms by which they operate in this key process.

*Drosophila* neural stem cells or NBs have been used as a paradigm for many decades to study ACD (51). NBs divide asymmetrically to give rise to another self-renewing NB and a daughter cell that will start a differentiation process. Over all these years, a complex network of ACD regulators that tightly modulate this process has been characterized. For example, the so-called cell-fate determinants, including the Notch inhibitor Numb, accumulate asymmetrically at the basal pole of mitotic NBs and are exclusively segregated to one daughter cell, promoting in this cell a differentiation process. The asymmetric distribution of cell-fate determinants in the NB is, in turn, regulated by an intricate group of proteins asymmetrically located at the apical pole of mitotic NBs, generically known as the “apical complex”. This apical complex includes kinases (i.e. aPKC), small GTPases (i.e. Cdc42, Rap1) and Par proteins (i.e. Par-6, Par3), among others (51).

Given the potential relevance of ACD in CSCs, we decided to take advantage of all the knowledge accumulated in *Drosophila* about the network of modulators that control asymmetric NB division. As a first approach, we aimed to analyze whether the levels of human homologs of known *Drosophila* ACD regulators were altered in human tumors. Specifically, we centered on human GBM, as the presence of CSCs (GSCs) have been shown in this tumor. The microarray we interrogated with GBM patient samples had some limitations. For example, not all the human genes homologs of the *Drosophila* ACD regulators were present (i.e. the human homologs of the determinant Numb). Likewise, we only tested seven different GBM patient samples. Nevertheless, the output from this analysis was enough to determine that most of the human genes tested in the array presented altered levels of expression. We selected for further analyses *RAP2A*, one of the human genes that showed the lowest levels of expression, compared to the control samples. However, it would be interesting to analyze in the future the potential consequences that altered levels of expression of the other human homologs in the array can have in the behavior of the GSCs. In silico analyses, taking advantage of the existence of established datasets, such as the TCGA, can help to more robustly assess, in a bigger sample size, the relevance of those human genes expression levels in GBM progression, as we observed for the gene *RAP2A*.

We have previously shown that *Drosophil*a Rap1 acts as a novel NB ACD regulator in a complex with other small GTPases and the apical regulators Canoe (Cno), aPKC and Par-6 (32). Here we have shown that *Drosophila* Rap2l also regulates NB ACD by ensuring the correct localization of the ACD modulators Cno and Numb. We also have demonstrated that RAP2A, the human homolog of *Drosophila* Rap2l, behaves as an ACD regulator in GBM neurosphere cultures, and its restitution to these GBM cultures, in which is present at low levels, targets the stemness of GSCs increasing the number of ACDs. It would be of great interest in the future to determine the specific mechanism by which Rap2l/RAP2A is regulating this process. One possibility is that, as it occurs in the case of the *Drosophila* ACD regulator Rap1, Rap2l/RAP2A is physically interacting or in a complex with other relevant ACD modulators. Thus, this study supports the relevance of ACD in CSCs of human tumors to refrain the expansion of the tumor. Other studies, however, claim that ACD should be targeted in human tumors as it promotes the intratumoral heterogeneity that hampers the complete tumor loss after chemotherapy (52-54). More investigations should be carried out with other human genes homologs of ACD regulators to further confirm the results of this study. Likewise, analyses in vivo (i.e. in mouse xenografts) would be also required to reinforce our conclusions. This would be very relevant in order to consider ACD restitution in CSCs of human tumors, what has been called “differentiation therapy” (55), as a potential alternative therapeutic treatment.

## Materials and Methods

### Drosophila husbandry, strains and genetics

The following fly stocks were used: *wor-Gal4 ase-Gal80 (56)*; *UAS-CD8::GFP* (Bloomington *Drosophila* Stock Center -BDSC-#5137); *UAS-CD2* (BDSC #1373); *UAS-Rap2l*^*KKRNAi*^ (Vienna *Drosophila* Resource Center -VDRC-#107745); *UAS-Rap2l*^*GDRNAi*^ (VDRC #45228); *UAS-Rap2l*^*RNAi*^ (BDSC #51840); *cno::GFP (57)*. All the fly stocks were raised and kept in 18^°^C or 25^°^C incubators. Experimental temperatures for the assays were maintained using 25^°^C or 29^°^C incubators. The *Gal-4* x *UAS*-crosses were carried out at 29^°^C (the first 2 days they were kept at 25^°^C, and then transferred to a new tube and left at 29^°^C) until third instar larvae developed.

### Drosophila histology, immunofluorescence and microscopy

Larval brains were dissected out in PBS and fixed with 4% PFA in PBT (PBS and Triton X-100 0.1%) for 20 min at room temperature with gentle rocking. Fixed brains were washed 3 times for 15 min with PBT (PBS and Triton X-100 0.3%) and then incubated in PBT-BSA for at least 1h before incubation with the corresponding primary antibody/antibodies. The following primary antibodies were used in this study: guinea pig anti-Dpn (1:2,000) (57), rabbit anti-Ase (1:100) (57), rabbit anti PKCζ (1:100; Santa Cruz Biotechnology, sc-216), goat anti-Numb (1:200; Santa Cruz Biotechnology, sc-23579), mouse anti-PH3 (1:2,000; Millipore, 05-806), mouse anti-γTub (1:200; Sigma-Aldrich, T5326) and mouse anti-CD2 (1:50; BioRad). Secondary antibodies conjugated to fluorescent dyes (Invitrogen) were used at 1:400 dilution. Samples were mounted in VECTASHIELD antifade mounting medium for fluorescence (Vector Laboratories, H-1000). Fluorescent images were captured using a Super-resolution Inverted Confocal Microscope Zeiss LSM 880-Airyscan Elyra PS.1. Images were analyzed using the image processing package FIJI from ImageJ and assembled using Adobe Photoshop CS6.

### GBM microarray

The cohort of patients consisted of seven GBM biopsies obtained from the Valencian Network of Biobanks. The procedures for obtaining tissue samples were developed in accordance with national ethical and legal standards, and following the guidelines established in the Declaration of Helsinki, and approved of Clinical Research Ethics Committee of the Hospital General Universitario de Elche, Spain. All patients had been diagnosed with GBM grade IV, and all of them exhibited an *IDH1* plus *IDH2* wild-type genotype. Normal human brain RNA (frontal lobe; pool from 5 donors) was purchased from BioChain and used as a control for gene expression experiments. A total of 12,516 genes were used to generate a custom Agilent Two-Color 8 x 15 K Agilent gene expression DNA microarray, which were deposited in the NCBI Gene Expression Omnibus (GEO) Database under accession number GSE182697 (28).

### Bioinformatics analysis

The level of expression of human homologs of *Drosophila* ACD regulators was assessed in seven different GBM patients versus control samples of the custom-made microarray (GSE182697) (28). Statistical analysis of microarray data was done in Multiexperiment Viewer (MeV) (58). Hierarchical clustering analyses were done optimizing gene and sample leaf order and using Pearson Correlation for the distance metric selection. Average linkage clustering was used as a linkage method. For the Gene Distance Matrix (GDM), the distance, inverse of similarity, between two genes was calculated using a distance metric as indicated in MeV.

### Human samples and cell lines

Glioma samples were obtained with patients’ written informed consent, in accordance with the Declaration of Helsinki, and approved by the Ethics Committee at Hospital 12 de Octubre, Madrid, Spain (CEI 14/023). The primary glioblastoma cell lines were provided by the Biobank at “Hospital 12 de Octubre”. Fresh tissue samples were enzymatically dissociated using Accumax (Millipore) and cultured in stem cell medium (Neurobasal, Invitrogen), supplemented with N2 (1:100, Invitrogen), GlutaMAX (1:100, Invitrogen), penicillin-streptomycin (1:100, Lonza), 0.4% heparin (Sigma-Aldrich), 40 ng/ml EGF and 20 ng/ml bFGF2 (Peprotech). The GB cell lines used in this study were: GB4, GB5, GB7, GB18, GB19, 12o89, Astros (59,60) and HEK293T. The cell lines HEK293T and Astros were maintained in Dulbecco’s modified Eagle Medium supplemented with 10% fetal bovine serum 2 mM L-glutamine, 0.1% penicillin (100 U/ml) and streptomycin (100 μg/ml) and Non-Essential Aminoacids (NEA 1x).

#### GBM-derived neurosphere cell cultures

For stem cell culture of GBM-derived neuroespheres we used DMEM:F12 with FGF-2 (20 ng/ml), EGF-2 (20 ng/ml), and N2 supplements, 0.1% penicillin (100 U/ml) and streptomycin (100 μg/ml), as previously described (61,62). The tumor spheres were dissociated to single cells using Accumax, and 1×10^5^ dissociated cells/ml were plated in a Flask and cultured for 6-7 days.

### In Silico Analysis

Data from The Cancer Genome Atlas (TCGA) and the Chinese Glioma Genome Atlas (CGGA) for GBM cohorts were accessed through UCSC Xena-Browser (https://xenabrowser.net), and Gliovis (http://gliovis.bioinfo.cnio.es) for extraction of overall survival data, gene expression levels, and distribution of genetic alterations. Kaplan-Meier survival curves were generated by stratifying samples into low- and high-expression groups for each gene. Survival differences between these groups were assessed using the log-rank test.

### DNA Constructs and Lentiviral Production

Lentiviral vectors were used to produce cells overexpressing *RAP2A* (*pLJM1RAP2A* #Plasmid 19311, Addgene) or eGFP (pLJMEGFP #Plasmid 19319, Addgene) as infection control. Infected cells were selected with Puromycin. To obtain the lentivirus, HEK293T cells were transiently co-transfected with 5 μg of the appropriate lentiviral plasmid (pLJEGFP or pLJRRap2A, respectively), 5 μg of the packaging plasmid pCMVdR8.74 (#Plasmid 22036, Addgene), and 5 μg of VERSUSV-G envelope protein plasmid pMD2G (#Plasmid 12259, Addgene), using Lipofectamine Plus reagent (Invitrogen, Carisbad, CA, USA). Lentiviral supernatants were collected after 48 h of HEK293T transfection.

### Neurosphere Immunofluorescences and Imaging

For immunostaining, floating cultured neurospheres were incubated on Matrigel coated micro slide glass for 2 hours at 37°C. After having been washed, the attached neurospheres were fixed in 4% paraformaldehyde for 20 min. Cells were blocked for 1 h in 1% FBS and 0.1% Triton X-100 in PBS; then, they were incubated for 2 hours with the primary antibody: mouse anti-CD133 (HB#7, Developmental Studies Hybridoma Bank), mouse anti-Ki67 (3E6, Developmental Studies Hybridoma Bank), rabbit anti-Sox2 (N1C3, GTX101507 GeneTex), mouse anti-Nestin (10c2, sc-23927, Santa Cruz Biotechnology) and mouse anti-Numb (48, sc-136554, Santa Cruz Biotechnology). The secondary antibodies used were Alexa555 goat anti-rabbit (A32732, Invitrogen) or Alexa555 goat anti-mouse (A32727, Invitrogen). Phalloidin 633 (Sigma-Aldrich, 68825 Phalloidin-Atto 633) was added and incubated for 20 min in PBS; Vectashield Mounting medium with DAPI (Linaris, H-1200) was added before mounting. Fluorescent images were captured using a Super-resolution Inverted Confocal Microscope Zeiss LSM 880-Airyscan Elyra PS.1. Images were analyzed using the image processing package FIJI from ImageJ and assembled using Adobe Photoshop CS6 program.

#### Immunofluorescent quantification

Fluorescence intensity measurements were carried out using ImageJ software package. In all cases, images of neuroespheres transfected with GFP (GB5, control) or RAP2A (GB5-RAP2A) lentivirus were analyzed:

<u>CD133:</u> Using the “line tool”, a line was drawn on the channel with the CD133 signal on three different pictures per genotype. Fluorescence was measured across the line sections with the “Plot Profile Tool”, and reported as a subset of images showing the signal intensity plot. For quantification, only the 100 highest gray value points, were taken.

<u>Sox:</u> After adjusting the threshold on the channel with the Sox2 signal and eliminating particles smaller than 5 microns, the fluorescence intensity was measured as the mean gray value (sum of all pixels with gray values inside the selection, divided by the number of pixels). In Figure 3 is shown one example out of three quantifications achieved with at least 20 cells per genotype.

<u>Nestin:</u> The regions of interest (ROIs) were set with the “hand drawing tool” using the Phalloidin staining to follow the cell periphery. At least 20 cells (ROIs) from three different pictures were selected per condition. The background fluorescence was subtracted from the mean gray value fluorescence for each well, and the fluorescence intensity was normalized by the nuclear DAPI signal coming from the same cell (ROI). Plot in Figure 3C represents the mean gray value that is the sum of all pixels with gray values inside the selection divided by the number of pixels.

### RNA isolation and qRT-PCR assay

Total RNA was extracted from 7 days old neurospheres using TRI reagent (AM9738, Invitrogen) following manufacturer indications and quantified using a nanodrop (ND-1000, Thermo Scientific). 2ug of RNA was treated with DNaseI (EN0521, Thermo Scientific) and reverse transcribed with SuperScriptTM III Reverse Transcriptase (1808044, Invitrogen). Quantitative real time PCR (qRT-PCR) was performed using Power SYBR Green PCR Master Mix (PN4367218, Applied Biosystems), following established protocols with 60^°^C for annealing/extension and 40 cycles of amplification on a QuantStudioTM 3 apparatus (applied Biosystems). Act5 primers were used for normalization, and relative expression was calculated using the comparative ΔΔCt. The average of at least 3 independent experiments is shown in Figure 4.

### Neurosphere Western Blots

Protein extracts were prepared by re-suspending cell pellets in a lysis buffer (50-mM Tris, pH 7.5, 300-mM NaCl, 0.5% SDS, and 1% Triton X-100) and incubating the cells for 10 min at 95°C. The lysed extracts were centrifuged at 13,000 g for 10 min at room temperature, and the protein concentration was determined using a commercially available colorimetric assay (BCA Protein Assay Kit). Approximately 25 µg of protein was resolved by 10% or 12,5% SDS-PAGE, and they were then transferred to a PVDF membrane (Hybond-ECL, Amersham Biosciences). The membranes were blocked for 1 h at room temperature in TBS-T (10-mM Tris–HCl, pH 7.5, 100-mM NaCl, and 0.1% Tween-20) with 5% skimmed milk, and then incubated overnight at 4oC with the corresponding primary antibody diluted in TBS-T with 2,5% skimmed milk. After washing 3 times with TBS-T, the membranes were incubated for 2 h at room temperature with their corresponding secondary antibody, diluted in TBS-T with 2,5% skimmed milk. The proteins were visible by enhanced chemiluminescence with ECL (Pierce) using Amersham imager 680, and the signal was quantified by Fiji-ImageJ software.

### Proliferative index (Ki67 analysis)

The number of Ki67+ cells with respect to the total number of nuclei labelled with DAPI within a given neurosphere were counted to calculate the Proliferative Index (PI), which was expressed as the % of Ki67+ cells over total DAPI+ cells. More than 20 neurospheres coming from different pictures were analyzed.

### Sphere size measurement

Glioblastoma cells (GB5 cell line) were infected by control lentivirus (LV-GFP) or by lentivirus directing expression of *RAP2A* (LV-*RAP2A*). The tumor spheres were Accumax-dissociated to single cells, during three consecutive passages. Seven days after the third plating in a 6-well plate, spheres were captured using a 10x objective (Carl Zeiss Axio Imager A1 microscope). 20 random pictures of each genotype were taken and the images were analyzed by measuring the area of all the spheres present, using the “freeform drawing tool” in the ImageJ software package. The data presented show the average area measured in 3 independent experiments.

### Pair assay and Numb segregation analysis

Dissociated cells were plated at low density (50.000 cells/ml) on Matrigel-coated well glass chamber slides on DMEM: F12 with FGF-2 (20 ng/ml), EGF-2 (20 ng/ml) and N2 supplements, 0.1% penicillin (100 U/ml) and streptomycin (100 μg/ml), supplemented with Puromycin. Cultures were incubated for 24, 48 or 72 hours at 37oC with 5% CO_2_. Bright field microscopy images from each time point were taken and then the slides were fixed and immunolabeled for Numb, as well as for DAPI to identify nuclei. First, isolated groups of cells (most probably coming from the same single progenitor) were identified and the number of cells in each cluster was annotated. Groups of 3-5 cells were counted in the slide incubated for 48h and groups of 6-8 cells in that incubated for 72h. We did not include in the analysis “neurospheres” with 2 cells, as these can be the result of a symmetric or an asymmetric cell division. We took into account neurospheres with 3-5-7 cells (as odd number of cells) and 4-6-8 cells (as even number of cells).

For the Numb analysis, the localization of Numb was visualized and scored in at least 100 DAPI-labeled cell pairs. To avoid any artefact, pairs were only captured in the slide incubated for 24h. Images in Figure 6 were acquired in an Inverted Microscope Leica Thunder Imager.

### Statistical analysis

The data were first analyzed using the Shapiro-Wilk test to determine whether the sample followed a normal distribution. Parametric T-test or a nonparametric two-tailed Mann Whitney U test for those that did not follow a normal distribution were used to compare statistical differences between two different groups. Mean and standard deviation values were calculated by standard methods. A Chi-square test was used to analyze the data in Figure 2C. To assume statistical significance, *p*-values were determined below 0.05. Sample size (n) and the *p*-value are indicated in the figure or figure legend; * *p* < 0.05, ** *p* < 0.01, *** *p* < 0.001, **** *p* < 0.0001; ns: not significant (*p* > 0.05).

## Acknowledgements

We thank the Bloomington *Drosophila* Stock Center at the University of Indiana and the Developmental Studies Hybridoma Bank at the University of Iowa for kindly providing fly strains and reagents. R.G. was funded by “Instituto de Salud Carlos III (ISCIII)” through the project “Miguel Servet Contracts” (CP21/00116), PI22/01171 and co-funded by the European Union. M.S. was supported by the Spanish grant from the Instituto de Salud Carlos III (ISCIII) PI22/00824. A.C. was financed by the Spanish grants from the Ministry of Science, Innovation and Universities (PGC2018-097093-B-100), the Ministry of Science and Innovation (PID2021-123196NB-100), the Generalitat Valenciana Prometeo/2021/052 grant and by FEDER (European Regional Development Fund). The “Instituto de Neurociencias” in Alicante is a “Severo Ochoa Center of Excellence in R&D”. Our group belongs to “Conexión Cáncer-CSIC”.

## Author contributions

M.F. participated in the design of the experiments, conducted most of them and analyzed the data; D.B. completed some experiments; R.G. developed the GB lines and performed the in silico analyses; V.M.B. and M.S. carried out the GB microarray and bioinformatics analysis; A.C. conceived the study and wrote the manuscript.

---

**Fig S1. Statistical analysis of asymmetric cell division expression levels in GBM patient samples**. A hierarchical clustering of genes showing significant or non-significant changes in their expression levels is shown. A t-test was applied to the microarray values of the asymmetric cell division regulator genes shown in Fig.1. The statistical values were presented in separate tables corresponding to the significant (**A**) and non-significant (**B**) changes in the indicated genes. The overall threshold *p*-value was 0.01. *p*-values were based on t distribution. Significance was determined using just Alpha.

**Figure S2. (A)** RAP2A expression levels analysis according to IDHwt GBM subtypes in the TCGA cohort. **(B-D)** Kaplan-Meier survival curves for each GBM subtype, Proneural **(B)**, Mesenchymal **(C)**, and Classical **(D)**, based on high and low RAP2A expression levels.

**Figure S3. *Drosophila RAP2A* homolog *Rap2l* regulates ACD**. Confocal micrographs showing a control larval brain NBII lineage with one NB (Dpn^+^ Ase^-^) and NBII lineages in which *Rap2l* has been downregulated displaying an ectopic NB (eNB). The phenotype of three different, independent *Rap2l*^*RNAi*^ lines (KK, GD and BDSC) are shown. Data represented in the bar graph was analyzed with a U Mann-Whitney test; \*\*\*\**p<0*.*0001*; n=number of NB lineages analyzed; the number of different brains analyzed per genotype is also indicated in brackets.

