## Supplementary figures and images for "Human *RAP2A* Homolog of the *Drosophila* Asymmetric Cell Division Regulator *Rap2l* Targets the Stemness of Glioblastoma Stem Cells"

### Supplemental Figure 1

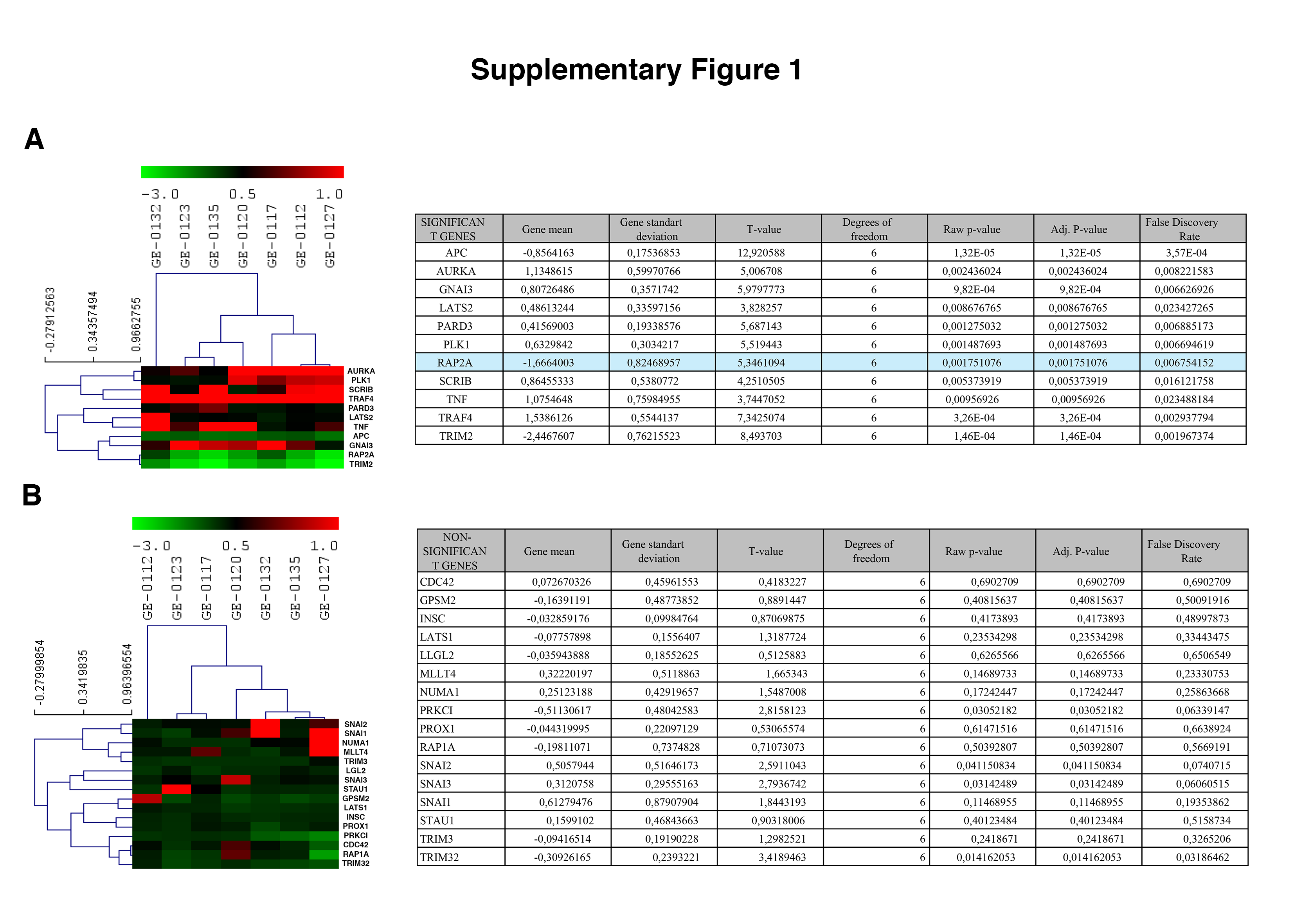

### Supplemental Figure 2

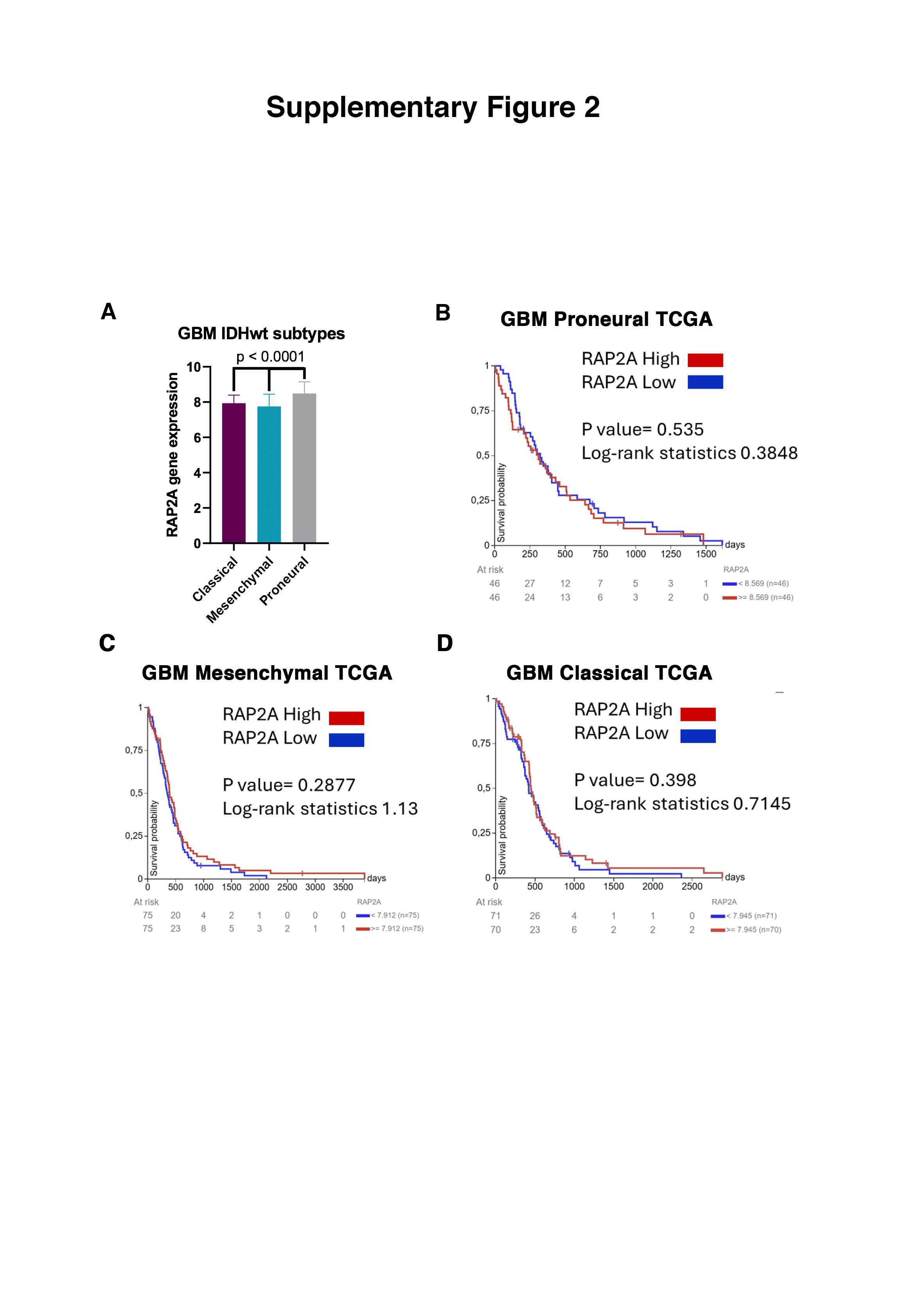

### Supplemental Figure 3

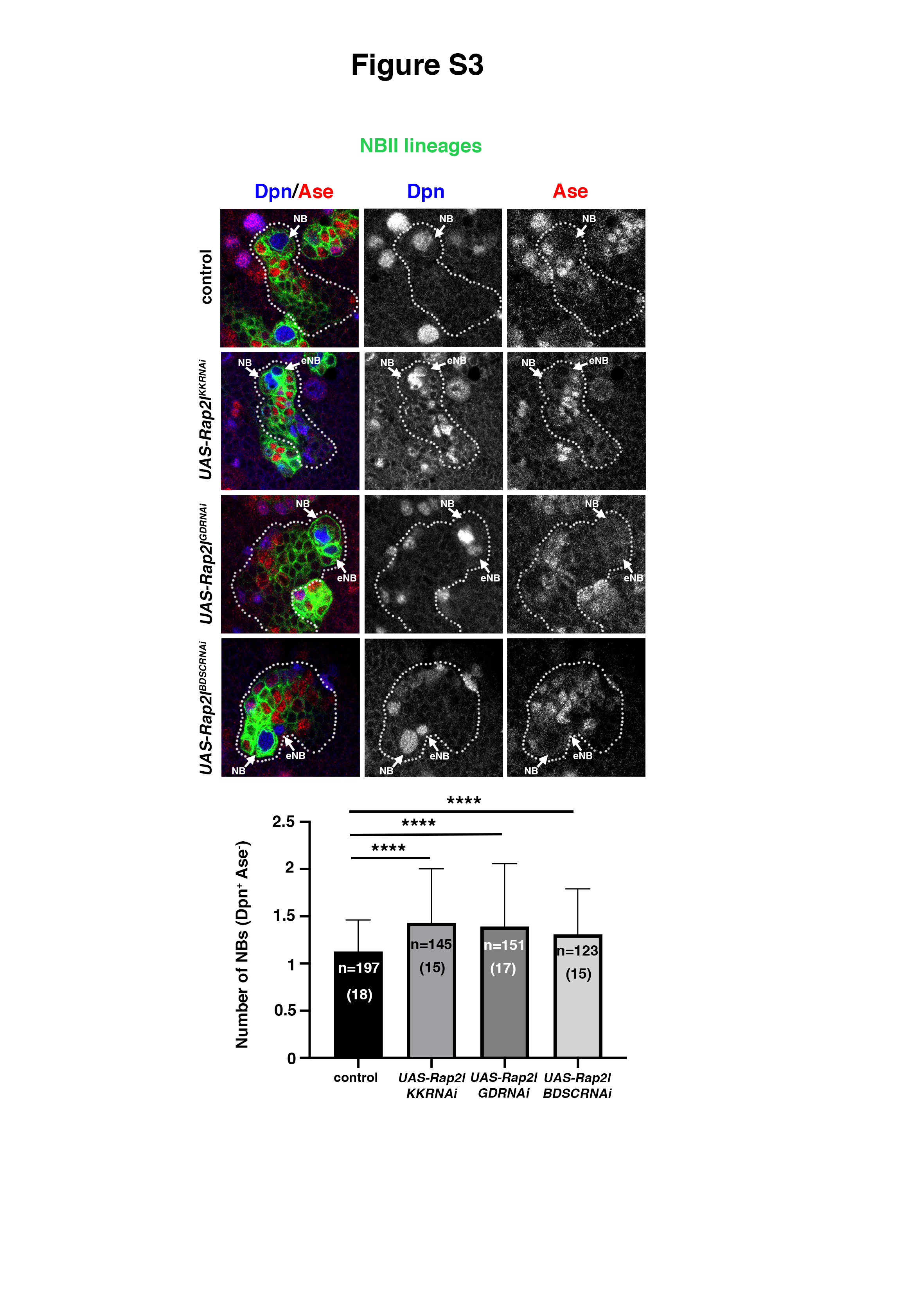
